# Preterm birth is associated with xenobiotics and predicted by the vaginal metabolome

**DOI:** 10.1101/2021.06.14.448190

**Authors:** William F. Kindschuh, Federico Baldini, Martin C. Liu, Kristin D. Gerson, Jingqiu Liao, Harry H. Lee, Lauren Anton, Pawel Gajer, Jacques Ravel, Maayan Levy, Michal A. Elovitz, Tal Korem

**Author notes:** These authors contributed equally to this work (William F. Kindschuh, Federico Baldini, Martin C. Liu).

## Abstract

Spontaneous preterm birth (sPTB) is a leading cause of maternal and neonatal morbidity and mortality, yet both its prevention and early risk stratification are limited. The vaginal microbiome has been associated with PTB risk, possibly via metabolic or other interactions with its host. Here, we performed untargeted metabolomics on 232 vaginal samples, in which we have previously profiled the microbiota using 16S rRNA gene sequencing. Samples were collected at 20-24 weeks of gestation from women with singleton pregnancies, of which 80 delivered spontaneously before 37 weeks of gestation. We find that the vaginal metabolome correlates with the microbiome and separates into six clusters, three of which are associated with spontaneous preterm birth (sPTB) in Black women. Furthermore, while we identify five metabolites that associate with sPTB, another five associate with sPTB only when stratifying by race. We identify multiple microbial correlations with metabolites associated with sPTB, including intriguing correlations between vaginal bacteria that are considered sub-optimal and metabolites that were enriched in women who delivered at term. We propose that several sPTB-associated metabolites may be exogenous, and investigate another using metabolic models. Notably, we use machine learning models to predict sPTB risk using metabolite levels, weeks to months in advance, with high accuracy. We show that these predictions are more accurate than microbiome-based and maternal covariates-based models. Altogether, our results demonstrate the potential of vaginal metabolites as early biomarkers of sPTB and highlight exogenous exposures as potential risk factors for prematurity.

## Introduction

Preterm birth (PTB), childbirth before 37 weeks of gestation, is the leading cause of neonatal death, and may lead to a variety of gastrointestinal, neurological, and other lifelong morbidities^1,2^. Despite extensive medical and research efforts to prevent PTB and ameliorate its consequences, its prevalence remains high both globally and specifically in the United States^1^. PTB also reflects a significant racial disparity, manifesting in a substantially higher risk for PTB in Black women compared to non-Hispanic white women^3^. Spontaneous preterm birth (sPTB), preterm birth not arising due to medical indication, accounts for two thirds of all PTBs, yet despite extensive study, methods for early prediction, prevention or treatment of sPTB are lacking^1,4–6^.

The human microbiome is a strong biomarker of many complex diseases^7–11^, often predicting host phenotypes even better than host genetics^12^. The vaginal microbiome, specifically, is a promising area of research for early diagnosis of sPTB: studies in multiple cohorts, clinical settings, and populations have found it to be associated with sPTB and other adverse pregnancy outcomes^13–18^. However, while multiple associations have been shown, a clear consensus on the relationship between the vaginal microbiome and sPTB has yet to emerge^19^. Several studies of the vaginal microbiome have reported associations between sPTB and microbial diversity or microbiome community state types (CSTs), but many of these associations have not generalized across cohorts^14,15,17–19^. Some of these discrepancies may be linked to the underlying structure of the population studied. For example, Callahan et al.^14^ found lower *Lactobacillus* and higher *Gardnerella* abundances to be associated with sPTB risk in a low risk, predominately white cohort, but not in a high risk, predominantly African American cohort^14^. In addition, our knowledge of specific mechanisms underlying potential host-microbiome interactions in sPTB is lacking.

Metabolites produced or modified by the microbiome have emerged as a prominent factor with potential local and systemic effects on the host^20–23^. Metabolomics enables the measurement of thousands of small molecules present in an ecosystem, and has revealed intriguing new insights on host-microbiome and other interactions in a variety of contexts, from colorectal cancer^24^ to diabetes^25^. A few studies of the vaginal metabolome have shown that it is associated with sPTB^26,27^. However, they had a limited sample size, and were not paired with measurements of the microbiome. Such paired microbiome-metabolome studies have yielded potential mechanistic insights in other pathologies, including inflammatory bowel disease^28^ and HPV infection^29^. The associations reported between both the vaginal microbiome and sPTB and the metabolome and sPTB highlight the need for paired analysis in order to advance our understanding of the role of this ecosystem in pregnancy outcomes.

Here, we measured the levels of 748 metabolites from vaginal samples collected in the second trimester of 232 pregnant women, for whom the composition of the microbiota was previously characterized using 16S rRNA gene amplicon sequencing^15^. We show that the vaginal metabolome partially corresponds to CSTs, and reveal novel associations between metabolites measured early in pregnancy and subsequent sPTB. Exploring the potential origins of some of these metabolites, we propose that some are of an exogenous source, suggesting a novel risk factor and potential targets for novel prevention strategies. Finally, we demonstrate that the vaginal metabolome, measured early in pregnancy, can accurately predict subsequent preterm birth in held-out samples, and show that it is more accurate than prediction based on microbiome or clinical data. Our results demonstrate a promising new approach for studying potential causes of prematurity as well as for early risk stratification, and highlight the need to study environmental exposures as a potential factor in sPTB.

## Results

### Vaginal microbiota and metabolome from a large pregnancy cohort

We used mass spectrometry to profile 232 vaginal samples collected between 20-24 weeks of gestation from women with singleton pregnancies (Methods). The microbiota of these women was previously characterized from the same double shaft swab used in this study^15^. We included all women with available samples who had a subsequent spontaneous preterm delivery in the parent cohort^15^ (sPTB; N = 80), as well as similar controls who delivered at term (TB; N = 152; **Table 1**). As expected, a higher fraction of women who delivered preterm had a history of PTB compared to those who delivered at term (42.5% vs. 19.2%, respectively, Fisher’s exact *p* = 3×10^−4^).

**Table 1:** Cohort characteristics. sPTB, spontaneous preterm birth; TB, term birth; BMI, body mass index; GA, gestational age; *p* - Fisher’s Exact or Mann-Whitney *U* test.

|  | sPTB | TB | Difference<br>(*p* value) |
| --- | --- | --- | --- |
| N | 80 | 152 |  |
| Race [N (%)] |  |  | 0.417 |
| Black | 57 (71.25%) | 116 (76.3%) | 0.568 |
| White | 21 (26.25%) | 30 (19.7%) | 0.331 |
| Other | 2 (2.5%) | 6 (4%) | 0.666 |
| Nulliparous [N (%)] | 29.0 (36.2%) | 55.0 (36.2%) | 0.894 |
| PTB history [N (%)] | 34 (42.5%) | 28 (19.2%) | **0.0003** |
| GA at delivery [median weeks [range]] | 34 [21-36] | 39 [38-39] | **<1e-10** |
| BMI [kg/m<sup>2</sup> mean±SD] | 30.1±7.8 | 30.6±7.2 | 0.65 |
| Age [years mean±SD] | 29±6 | 28±6 | 0.28 |

We quantified 748 unique metabolites, of which 637 could be named (Methods). Metabolites belonged to diverse biochemical classes, including amino acids, lipids, nucleotides, carbohydrates and xenobiotics. Most metabolites (549) were measured in over 50% of the cohort, and 108 metabolites were present in all samples (**Fig. S1**). We have previously shown that similar metabolite measurements are in excellent agreement with measurements performed by an independent certified medical laboratory^30^.

### The vaginal metabolome partially preserves CST structure

The vaginal microbiome clusters to well-defined community state types (CSTs)^31^. We demonstrated the same for this cohort^15^ (PERMANOVA *p* < 0.001 for separation between CSTs; **Fig. 1a**), and investigated whether the vaginal metabolome space recapitulates this structure. While the metabolome is separated by CSTs (*p* < 0.001; **Fig 1b**), and is generally associated with the microbiome (Mantel *p* < 0.001), specific CSTs are not as well separated. While women with microbiomes characterized as CST I, dominated by *Lactobacillus crispatus*, and CST IV, characterized by diverse anaerobes, are well separated from the rest of the cohort in their metabolite measurements (PERMANOVA *p* < 0.001 for both), neither the metabolite measurements from women with CSTs IV-A and IV-B, nor CSTs II and III, were well separated from one another (*p* = 0.169 and *p* = 0.171, respectively). Overall, these results demonstrate a strong but imperfect correspondence between the vaginal microbiome and metabolome spaces.

**Figure 1.**
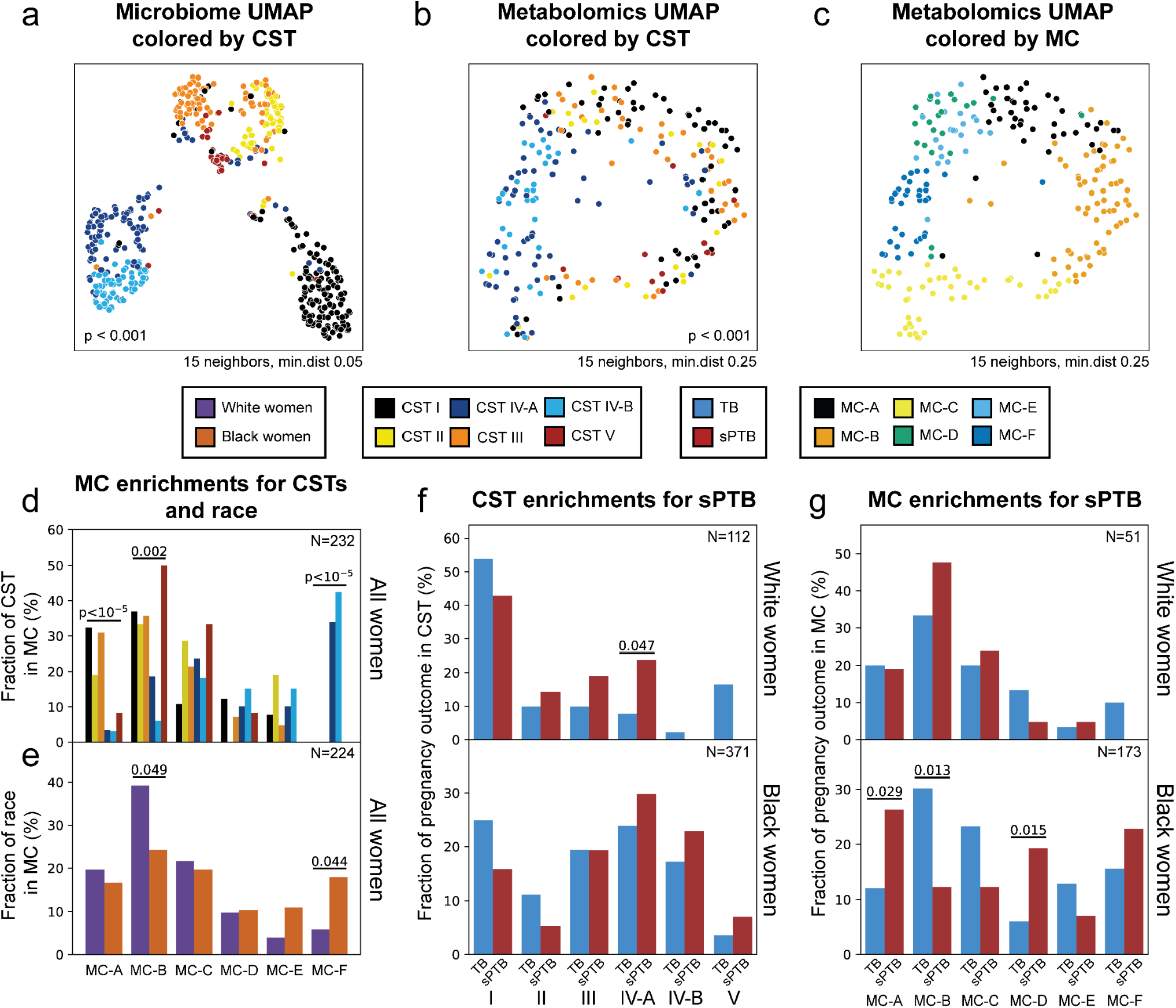
Vaginal metabolome clusters are associated with preterm birth. **a-c**, UMAP ordination of microbiome (a; N = 503) and metabolomics data (b,c; N = 232), colored by community state types (CSTs; a,b) or de-novo clustering of metabolites data (metabolites clusters [MCs]; Methods; c). The vaginal microbiome and metabolome are significantly separated by CSTs (PERMANOVA *p* < 0.001 for both), yet the separation is less clear in the metabolome. **d**, The fraction of women whose metabolite profiles clustered to each MC, shown for each CST separately. **e**, Similar to **d** but shown for Black and white women separately. **f**, The fraction of white (top) and Black (bottom) women whose microbiomes belonged to each CST, separated by pregnancy outcome. As previously shown^15^, CST IV-A is enriched for sPTB among white women (*p* = 0.047). **g**, Similarly as **f**, for the fraction of women whose metabolomes clustered to each MC. We show a significant association of sPTB with MC-A, B and D among Black women (*p* = 0.029, *p* = 0.013, *p* = 0.015, respectively). *p* - Fisher’s exact.

### Metabolome-clusters associate with sPTB

While the metabolome is significantly clustered to CSTs, this clustering is not perfect. We therefore performed de novo clustering of the metabolome using k-medoids clustering, and revealed six “metabolite-clusters” (MCs A-F; Methods; **Fig. 1c, S2**,**3**; **Table S1**). We find that amino-acid-related metabolites are overrepresented among metabolites significantly associated with MC-A, MC-B, and MC-D (Mann-Whitney *U p* < 0.05) compared to other MCs (Fisher’s exact *p* = 4.3×10^−8^, *p* = 0.0011, *p* = 1.8×10^−8^, respectively; FDR < 0.1 for all) and that xenobiotics are overrepresented among metabolites associated with MC-C (Fisher’s exact *p* = 0.0014, FDR < 0.1). We further show that while three MCs are mostly paired with *Lactobacillus* dominated CSTs (57%-93% for MC A-D), MC-F is composed entirely of CST IV, and MC-D and E are evenly split (50-52% CST IV; **Fig. 1d, S4a**). Reciprocally, we found various enrichments of CSTs in MCs (**Fig. 1d, S4b)** as well as enrichments for white women in MC-B and Black women in MC-F (*p* = 0.049 and *p* = 0.044, respectively; **Fig. 1e**). These results demonstrate that the correspondence between the vaginal metabolome and microbiome is imperfect, and somewhat corresponds to *Lactobacillus* dominance.

A previous analysis of the same cohort showed that CST IV-A is associated with subsequent sPTB only among white women^15^ (Fisher’s exact *p* = 0.047; **Fig. 1f, S4c**). We therefore performed the same stratified test using MCs. Interestingly, while CSTs are associated with sPTB only in white women, we find that MCs are associated with sPTB in Black women, with a significant association with MC-A, MC-B, and MC-D (*p* = 0.029, *p* = 0.013, *p* = 0.015, respectively; **Fig. 1g, S4d**). Taken together, our results demonstrate that the general metabolome structure in our cohort better captures associations with prematurity in Black women than the general microbiome structure.

### Multiple metabolites associate with sPTB

Having demonstrated that the global metabolome structure associates with sPTB, we next investigated its associations with the levels of specific metabolites. We find five metabolites that are significantly associated with sPTB (Mann-Whitney *U p* < 0.05, FDR < 0.1; **Fig. 2a**). Three of these, all higher in women who delivered preterm (*p* < 10^−3^, FDR < 0.1 for all; **Fig. 2a**), appear to be of exogenous source: ethyl glucoside (ethyl beta-glucopyranoside; *p* = 1.9×10^−4^), an alkyl glucoside used as a surfactant in cosmetic products^32^; tartrate (*p* = 4.8×10^−4^), used, along with its products, in the food, pharmaceutical, and cosmetics industries^33,34^; and diethanolamine (DEA; *p* < 10^−10^), commonly used in personal care and cosmetic products^35^, and which was shown, in mice, to cause liver and kidney tumors^36^, as well as induce apoptosis in fetal hippocampus cells, decrease neural progenitor cell mitosis in the hippocampus, and reduce litter size in a dose dependent manner^37^. We note, however, that due to the nature of our metabolomic measurements, the levels of these metabolites in the vaginal ecosystem may be different than those observed in previous studies.

**Figure 2.**
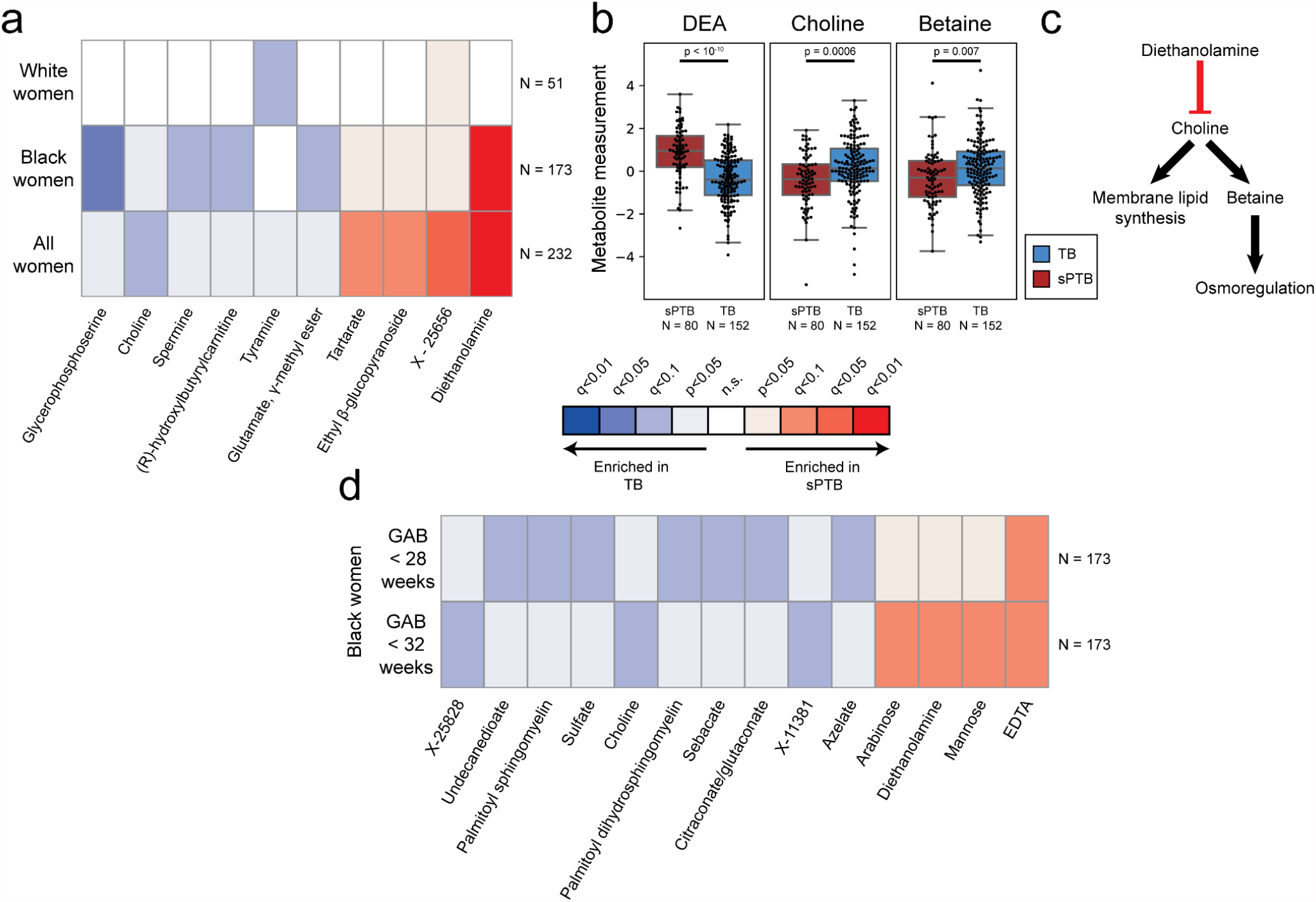
Vaginal metabolites associate with subsequent preterm delivery. **a**, Heatmap showing statistically significant associations (Mann-Whitney *U p* < 0.05) between specific metabolite measurements and birth outcomes, stratified by maternal race, and colored by significance and direction of association. Only metabolites with at least one association with FDR < 0.1 are shown. Metabolites are sorted by their average signed (direction of fold change) log *p*-value. **b**, Box and swarm plots (line, median; box, IQR; whiskers, 1.5*IQR) of three metabolites with significant associations with sPTB. *p* - Mann-Whitney *U*. **c**, Illustration summarizing some of the literature regarding the three metabolites shown in **b**. Diethanolamine (DEA), which is associated with sPTB, was shown to inhibit choline uptake^41^. Choline and betaine, both associated with TB, are important for membrane lipid synthesis and osmoregulation^38,40^. **d**, Same as **a**, with stratification by gestational age at birth (GAB), performed among Black women. Middle legend applies to **a** and **d**.

We further find lower levels of the amine choline in women with subsequent sPTB (*p* = 5.5×10^−4^, FDR < 0.1; **Fig. 2a,b**). Choline is an essential nutrient required for membrane phospholipids and neurotransmitter synthesis^38^, and lower choline levels were previously found in cord blood from premature infants^39^. Choline is also a precursor of betaine^40^, a metabolite mainly involved in osmoregulation^40^ which was also negatively associated with sPTB (*p* = 0.007, FDR = 0.14; **Fig. 2b**). DEA is known to disrupt several enzymes and transporters involved in choline metabolism^41^, and its dermal administration in mice was reported to deplete hepatic stores of choline^37,42^. We therefore propose that the high levels of the exogenous chemical DEA, which was higher in samples from women who delivered preterm, may be linked to the lower choline and betaine levels that we found (**Fig. 2b,c**). Taken together, these results highlight a potential role of exogenous metabolites in prematurity, potentially from environmental exposures via hygienic and cosmetic products.

### Metabolite associations with sPTB interact with race and sPTB timing

As the global metabolome structure shows differences between Black and white women, we performed the same association analysis while stratifying by race. Interestingly, we detect five additional metabolites negatively associated with sPTB (Mann–Whitney *U p* < 0.05; FDR < 0.1; **Fig 2a**). In Black women, these include glycerophosphoserine (*p* = 3×10^−5^), which was previously reported to be altered in preeclampsia^43^; spermine (*p* = 2.4×10^−4^), which has immuno-modulatory roles in the gut^23^, and was increased in the blood of preterm infants^44^; hydroxybutyl carnitine (*p* = 2.6×10^−4^), a ketocarnitine involved in lipid metabolism and which has been shown to be depleted in the blood of low birth weight full-term neonates^45,46^; and glutamate gamma-methyl ester (*p* = 4.9×10^−4^), a derivative of glutamate and a precursor of the inhibitory neurotransmitter GABA. Tyramine, a biogenic amine with neuromodulator activity^47^, was significantly lower in samples from white women who delivered preterm (*p* = 2.8×10^−4^; **Fig. 2a**). Tyramine was shown to colocalize with synaptic vesicles in the mouse uterine plexus, highlighting a possible role in uterine contractions^48,49^. Altogether, these results highlight the potential connection between vaginal metabolites, metabolite levels in various reproductive organs, and preterm delivery.

Earlier preterm deliveries are associated with worse maternal and neonatal outcomes^1^. Therefore, we next investigated associations between vaginal metabolites and subsequent very and extremely preterm deliveries (gestational age at birth <32 and <28 weeks, respectively). Due to the high proportion of Black women among pregnancies delivering prior to 32 and 28 weeks (21 of 26 and 14 of 15, respectively), we performed this analysis only among Black women. We identify 12 metabolites that are associated only with these earlier sPTBs (**Fig. 2d**). The phospholipids palmitoyl sphingomyelin and palmitoyl dihydro sphingomyelin were both negatively associated with extremely PTB (*p* = 0.00087 and *p* = 0.0011, respectively). Phospholipids, however, and specifically palmitoyl sphingomyelin, were recently found to be increased in placental tissue of sPTB deliveries^50^. Citraconate, a derivative of citric acid, was likewise negatively associated with extremely PTB (*p* = 0.0014), and was previously found to have significantly lower concentrations in placental mitochondria of women with severe preeclampsia^51^. Ethylenediaminetetraacetic acid (EDTA) was one of two metabolites increased in extremely and very PTB (*p* = 0.0008 and *p* = 1.5×10^−4^, respectively). EDTA is another metabolite that can be found in cosmetic products, where it acts as a chelating agent^52^. Potentially at different concentrations that those measured here, EDTA has also been shown to be cytotoxic in vaginal epithelial cells, in which it provokes an inflammatory response^52,53^, and is teratogenic, causing fetal gonadal dysgenesis in rats at non-maternotoxic doses^52,54,55^. We note that while EDTA is also present in the buffer in which our samples were collected, this is unlikely to explain the observed association. Overall, we find that the associations between metabolites and sPTB interact with both race and timing of sPTB, and detect an additional xenobiotic that is associated with sPTB.

### Functional metabolite sets are enriched with sPTB-associated metabolites

In addition to identifying specific metabolites with strong associations with sPTB, we next checked whether certain functional groups of metabolites (e.g. KEGG pathways^56^) are enriched for associations with sPTB, compared to all other metabolites, even if changes to any specific metabolite are small in scale (**Fig. S5**; Methods). We find significant sPTB-associated deviations in metabolites related to proline and arginine metabolism (*p* = 0.0018, FDR < 0.1; **Fig. S5**). This set includes spermine, which has been shown to be derived from proline^57^, and was lower in Black women that delivered preterm (**Fig. 2a**). Proline itself comprises about a quarter of the amino acid residues of collagen^58^, and is therefore integral to the extracellular matrix. It is also converted to arginine and metabolized to form polyamines, which are important for placental angiogenesis^59^, but have also been associated with CST IV^60^. Both disordered placental angiogenesis and extracellular matrix remodeling have been associated with preterm birth^61^, possibly reflecting these changes in proline and arginine metabolism.

Consistent with the association we found between tyramine levels and term births among white women (**Fig. 2a**), we find a global deviation of metabolites related to the endocrine system among white women (*p* = 0.0046, FDR < 0.1; **Fig. S5**). We further identify lipid-metabolism-related metabolites to be enriched for associations with early sPTB among Black women (*p* = 0.0021 and *p* = 0.0048 for very and extremely PTB, respectively, FDR < 0.1; **Fig. S5**), potentially related to other alterations in lipid metabolism reported in women who delivered preterm^62,63^. Notably, we identify a global enrichment of xenobiotic metabolites associated with sPTB among Black women (*p* = 0.006, FDR < 0.1; **Fig. S5**). This is consistent with our finding of exogenous metabolites associated with sPTB in this population, and could potentially be related to the higher burden of exogenous and environmental exposures in Black communities^64,65^, which have been identified as potential drivers of preterm birth^66–69^. In all, our analyses highlight multiple metabolites associated with sPTB, and that these associations interact with both race and sPTB severity.

### A network of microbe-metabolite associations in sPTB

To obtain further insights regarding potential microbial sources of metabolites altered in sPTB, we next investigated associations between the absolute abundances of various microbial species and metabolites associated with sPTB (**Fig. 2a**; Methods). Our results replicate several known associations, such as those between *Dialister* species or *Enterococcus faecalis* and tyramine^60,70^ (Spearman ρ > 0.54, *p* < 10^−5^ for all; **Fig. 3a, S6**), as well as evidence for choline metabolism genes in *Gardnerella vaginalis*^71^ and *Corynebacterium aurimucosum*^72^ (ρ = 0.34, *p* < 10^−6^ and ρ = 0.40, *p* = 0.0004, respectively). Additionally, higher concentrations of tyramine were previously found in bacterial vaginosis (BV)^60,73,74^, supporting many of the associations we find with bacteria that are also associated with BV (**Fig. 3a**). Finally, we observe a positive correlation between *C. aurimucosum* and spermine (ρ = 0.27, *p* = 0.02), and it has been shown that spermine and its precursor spermidine are the key polyamines in several *Corynebacterium* species^75,76^.

**Figure 3.**
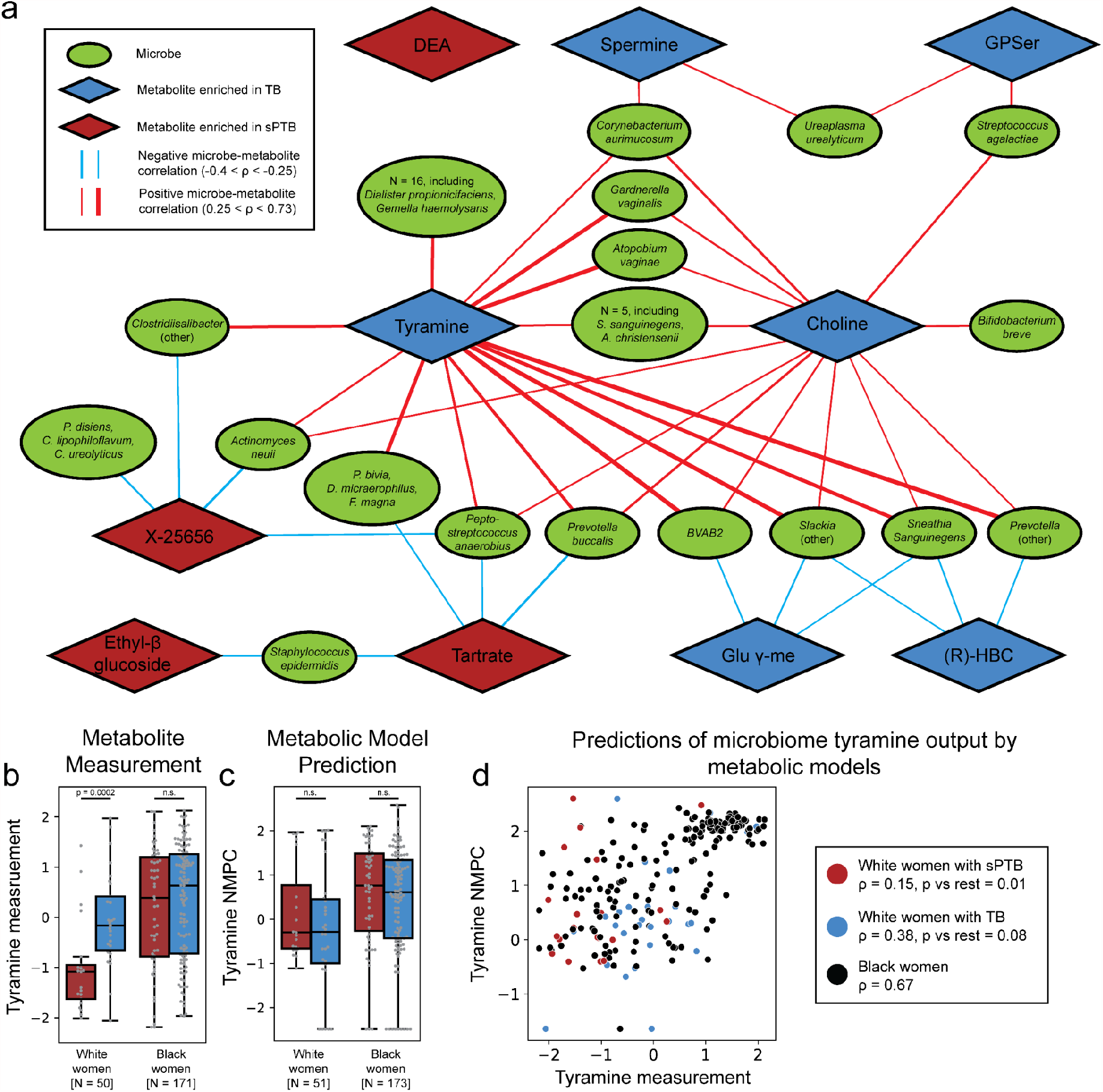
Microbe-metabolite correlations and metabolic models suggest sources for sPTB-associated metabolites. **a**, A network of microbial correlations with metabolites associated with sPTB. Ellipses, microbial species; blue and red diamonds, metabolites enriched in TB and sPTB, respectively; blue and red edges, negative and positive Spearman correlations with FDR < 0.05, |ρ| > 0.25, respectively (edge width corresponds to the median ρ). See **Fig. S6** for the same network without grouped nodes. **b**,**c**, Box and swarm plots (line, median; box, IQR; whiskers, 1.5*IQR) of tyramine levels, as measured (b) and predicted with metabolic models (Methods; c), comparing preterm and term deliveries and stratifying by maternal race. White women who delivered preterm had lower measured vaginal levels of tyramine (*p* = 0.0002), yet our metabolic models predict higher, albeit non-statistically significant, microbiome production of tyramine in women who delivered preterm (*p* = 0.16 and *p* = 0.24 for all and white women, respectively). *p*, Mann-Whitney *U*. **d**, Tyramine production derived from microbiome metabolic models (NMPC; Methods; y-axis) plotted against measured tyramine levels (x-axis) and colored by race and birth outcome (legend). While our models are generally accurate for tyramine (Spearman ρ = 0.63, *p* < 10^−10^ across all women), the accuracy for white women who delivered preterm was significantly lower (Spearman ρ = 0.15, *p* = 0.01 for comparing correlation strength vs the correlation in other women), suggesting a non-microbial interaction.

We find the strongest and most numerous microbe-metabolite correlations for tyramine (35 associations, Spearman 0.27 < ρ < 0.73; **Fig. 3a**) which was higher in term deliveries among white women (**Fig. 2a**). Eight out of the 35 tyramine-correlated microbes are also correlated with choline, which was enriched in term deliveries across all women (**Fig. 2a**). Interestingly, many of the species positively correlated with metabolites associated with term delivery, including *Atopobium vaginae, G. vaginalis, Sneathia sanguinegens, C. aurimucosum, Mobiluncus curtisii, Actinomyces neuii, Ureaplasma urealyticum, Gemella haemolysans*, several *Prevotella* species, *Candidatus Lachnocurva vaginae* (BVAB1^77^), BVAB2 and BVAB3 were previously reported to be associated with negative outcomes, such as BV^31,78–81^, preterm birth^14–16,18,82^ and other adverse pregnancy^82–84^ and neonatal^85^ outcomes. We find a similarly paradoxical negative correlation between *Staphylococcus epidermidis*, previously shown to be associated with BV^86^ and late-onset sepsis in preterm neonates^87^, and both tartrate and ethyl glucoside (ρ = -0.28, *p* = 0.00069; ρ = - 0.26, *p* = 0.0015, respectively; **Fig. 3a**), which were positively associated with sPTB. These results suggest the existence of a potential beneficial metabolic effect for some microbial populations that were so far considered dysbiotic.

As many of the associations between metabolites and sPTB were modulated by race, we next investigated whether the associations in our network are likewise influenced. We find that nine of the 75 microbe-metabolite associations we detected were significantly different (Fisher’s R-to-z *p* < 0.05; Methods) between Black and white women, although a different direction of association was detected in only four of these (**Fig S7**). Specifically, *G. vaginalis, A. vaginae*, and three other species that were positively associated with tyramine, had significantly stronger associations in Black women (*p* < 0.02 for all). Recent studies show that biofilm interactions between *G. vaginalis* and other microbes, including *A. vaginae*^88,89^, may contribute to BV, suggesting that the differences in tyramine associations between Black and white women may be related to differences in community structure and microbial interactions. Taken together, however, we find a relatively small effect of race on microbe-metabolite correlations.

### Microbiome metabolic models support microbial production of tyramine

To gain some mechanistic insight into the correlations we found, we next used community-level metabolic models^90^ to predict the metabolic output of each microbiome sample (community net maximal production capability^90^ [NMPC]; Methods). These models combine genetic and biochemical knowledge to generate predictions of metabolite output, using only microbial relative abundances^91^. We focused on tyramine, which was one of two sPTB-associated metabolites represented in our models (the other, choline, had no predicted microbial production) and which previous studies suggest is produced by vaginal microbes^60,74,92^. Following genomic curation of our metabolic models (Methods), the predictions of our models were highly accurate (Spearman ρ = 0.63 between tyramine NMPC and its metabolomic measurements, *p* < 10^−10^, N = 229 predictions). When examining samples from white women, we find that while the measured levels of tyramine were enriched in TB (Mann-Whitney *U p* = 0.00028; **Fig. 3b**), its predicted output by the microbiome was not, and was even somewhat higher in sPTB (*p* = 0.24; **Fig. 3c**). This stems from a lower accuracy in tyramine predictions in white women who delivered preterm (Spearman ρ = 0.15 versus ρ = 0.65, p = 0.01 for difference in ρ’s; **Fig. 3d**), suggesting a non-microbial effect on tyramine levels among white women and providing a potential explanation to the aforementioned paradoxical microbial associations with tyramine (**Fig. 3a**). Our results demonstrate the utility of metabolic models in potentially discerning microbial from non-microbial effects.

### Early prediction of sPTB risk using the vaginal metabolome

Early diagnosis of pregnancies with high risk for prematurity is crucial for the development of prevention and intervention strategies, yet is largely still lacking^1,4–6^. We therefore explored whether we can use our microbiome and metabolome data, collected at weeks 20-24 of gestation, to predict sPTB. Of note, prediction was of deliveries occurring up to 19 weeks after the samples were taken. We trained predictive models and evaluated them on held-out samples using multiple different draws of 10-fold cross-validation, with strict train-test sterility (Methods). As a benchmark, we constructed models using clinical data (age, BMI, race, parity status, history of sPTB and nulliparity), which obtains limited accuracy (auROC = 0.62, auPR = 0.45; **Fig. 4a,b**), similar to that obtained in past studies^4^. Using microbial abundances, we are able to only slightly improve prediction accuracy (auROC = 0.63, auPR = 0.49; *p* = 0.36 for comparison of auROC with clinical model; Methods; **Fig. 4a,b**).

**Figure 4.**
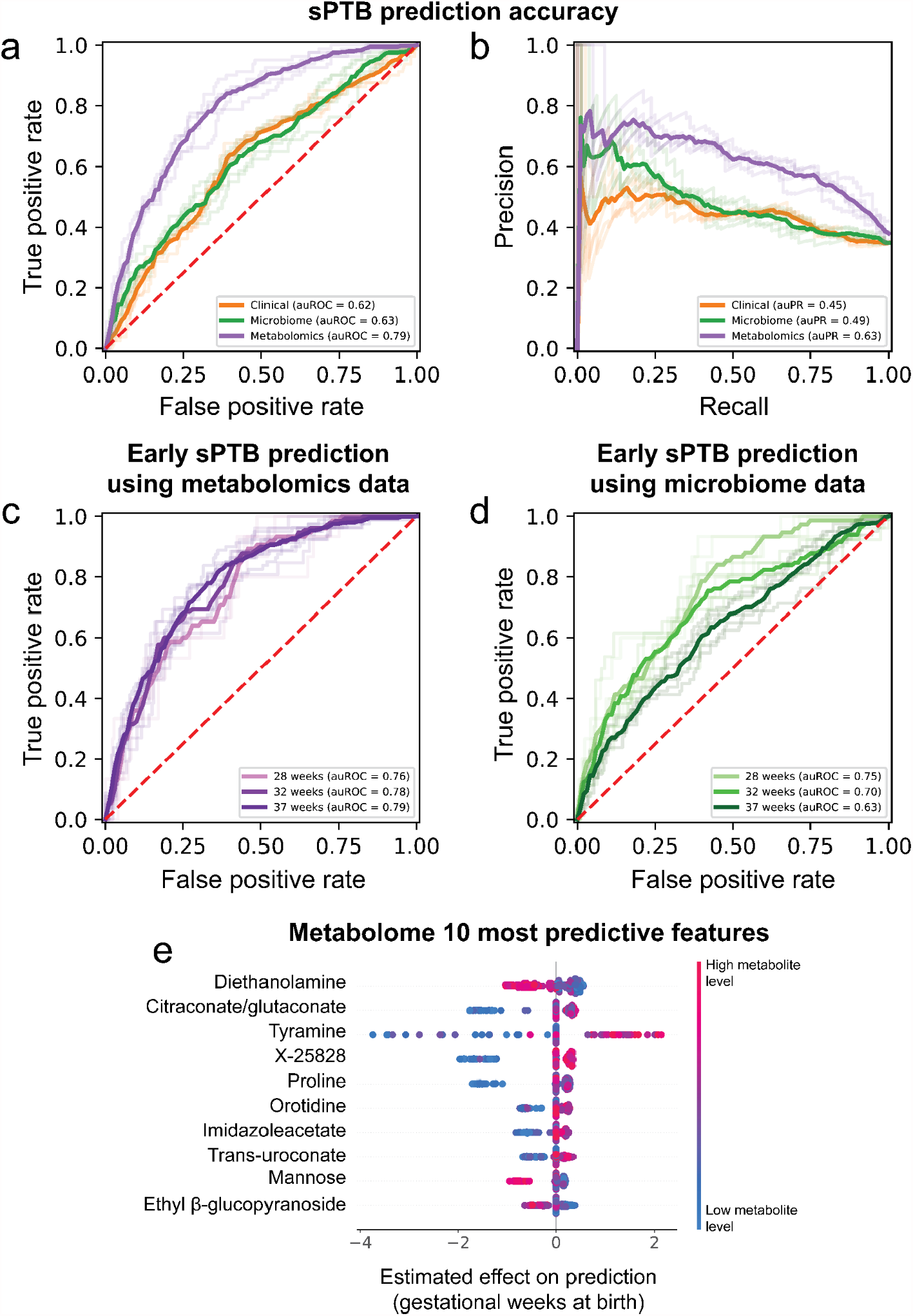
Metabolomics-based prediction of subsequent spontaneous preterm birth. **a**,**b**, Receiver operating characteristic (ROC, a) and Precision-recall (b) curves comparing sPTB prediction accuracy for models based on clinical (auROC = 0.62, auPR = 0.45), microbiome (auROC = 0.63, auPR = 0.49) and metabolomics (auROC = 0.79, auPR = 0.63) data. N = 232 for all. Shaded lines show results from five independent and random 10-fold cross-validation draws (Methods). **c**,**d**, Same as **a**, for early sPTB prediction based on metabolomics (c) and microbiome (d) data. Microbiome data shows significant improvement in predicting extremely PTB (<28 weeks of gestation; auROC of 0.75 compared to 0.70 and 0.63 for very and all sPTB, respectively), while metabolomics data show similar performance across sPTB severity (auROC 0.76-0.79 for all). **e**, Effect on total prediction (SHAP-based^98^; x-axis) for the 10 most predictive metabolites in our metabolomics-based predictor, sorted with descending importance. Each dot represents a specific sample, with the color corresponding to the relative level of the metabolite in the sample compared to all other samples.

Notably, using metabolomics data, we were able to generate a model with superior accuracy (auROC = 0.79; auPR = 0.63; *p* < 10^−10^ and *p* = 2×10^−8^ for comparison of auROCs with clinical and microbiome models, respectively; Methods; **Fig. 4a,b**). Lastly, a model combining clinical, microbiome, and metabolomics data obtains similar accuracy to the model which used only metabolomics data (auROC = 0.77, auPR = 0.63; *p* = 0.24 for comparison of auROC with metabolomics model; **Fig. S8a,b**), with metabolomics-based features as the most prominent contributors to the model (9 of 10 most predictive features; **Fig. S8c**). This suggests that metabolite measurements are a sufficient representation of information contained in these three data types. Our metabolomics-based model is superior or similar in accuracy to several previously-published models, such as those using amniotic fluid metabolomics (auROC = 0.65-0.70, N = 24)^93^, maternal serum metabolome and clinical data (auROC = 0.73, N = 164)^94^, maternal urine and plasma metabolomics (auROC = 0.69-0.79, N = 146)^95^, cell-free blood RNA measurements (auROC = 0.81, N = 38)^96^, or vaginal protein biomarkers (auROC = 0.86, N = 150, sPTB N = 11)^97^, many of which have small sample sizes, lack demographic diversity, or focus on high-risk cohorts. Overall, our results demonstrate the promising utility of vaginal metabolites as early and accurate biomarkers of spontaneous preterm birth.

We next checked whether microbiome- and metabolomics-based predictors are more capable at predicting early sPTB, which is associated with higher morbidity and mortality^1^. We therefore evaluated the same models, without retraining, for predicting extremely (<28 weeks) and very (<32 weeks) PTB. While our metabolites-based model maintains similar accuracies across different sPTB timings (auROC of 0.76 and 0.78 for extremely and very PTB, respectively, compared to auROC of 0.79 for all sPTB, *p* > 0.20 for both; **Fig. 4c**), our microbiome-based model shows substantially higher accuracy for predicting extremely PTB (auROC of 0.75 and 0.70 for extremely and very sPTB, respectively, compared to auROC of 0.63 for all sPTB, *p* = 1.5×10^−4^ and *p* = 2.3×10^−3^, respectively; **Fig. 4d**). Interestingly, these results reflect the potentially increased involvement of the vaginal microbiome in earlier sPTBs^1,99^.

### Interpretation of predictive models reveals novel contributing features

To interpret our predictive models and obtain insights into the features they use, we performed feature attribution analysis (using SHAP^98^) allowing us to infer the contribution of each feature towards the final prediction for each sample (**Table S2**). As expected, six of the ten most predictive metabolites, namely diethanolamine, tyramine, ethyl-glucoside, citraconate/glutaconate, mannose, and the unidentified metabolite X-25858, were also identified in our association analysis, with a similar direction of association **(Fig. 2, 4e)**.

In addition to these previously identified associations, our metabolomics-based model also relies on proline, with lower proline levels contributing towards sPTB prediction (proline is the fifth strongest feature; **Fig. 4e**). Taken together with the significant enrichment that we observed in sPTB associations among metabolites related to arginine and proline metabolism (**Fig. S5**), the strong predictive power of proline further suggests a role for extracellular matrix remodeling and polyamine metabolism in sPTB. Lastly, our analysis indicates that low levels of imidazoleacetate, orotidine, and trans-urocanate also contributed to sPTB predictions (**Fig. 4e**). Urocanic acid was previously shown to be increased in amniotic fluid of women who delivered preterm^100^. Additionally, both trans-urocanate and imidazoleacetate are derivatives of the amino acid histidine, and were correlated with it in our cohort (ρ = 0.33, *p* = 7.5*10^−7^ and ρ = 0.37, *p* = 5.1*10^−9^, respectively). Histidine levels were previously found to be lower in both second trimester amniotic fluid of women who delivered preterm^101^ and in urine from preterm infants^102^.

A similar analysis, performed on our microbiome-based predictor, has also captured previously-detected associations between various vaginal microbes and sPTB^15^, including those of *Mobiluncus mulieris, S. sanguinegens*, and of *Lactobacillus, Streptococcus*, and *Dialister* species (**Fig. S8d**). We further identify a new association with *Finegoldia magna*, an anaerobic gram-positive species whose levels contributed to prediction of sPTB (**Fig. S9d**). This replicates results from a previous study, which showed that *F. magna* was more prevalent in women who delivered preterm^103^. These results highlight the interpretability of our models and their reliance on complex, non-linear interactions of metabolites with both sPTB and other features, enabling us to expose associations not detected by univariate analyses.

## Discussion

In this study, we measured the levels of 748 vaginal metabolites in a cohort of 232 pregnant women. We show that the vaginal metabolome largely separates by microbial community state types, but that de novo clustering of the metabolome reveals clusters enriched for sPTB among Black women. We further identify multiple metabolites that are associated with sPTB, with differences between Black and white women and between early and late preterm births. Our results highlight several exogenous metabolites with strong associations with sPTB, which we suggest to be the result of environmental exposures. Using microbe-metabolite associations, we uncover intriguing interactions between metabolites associated with term birth and potentially suboptimal microbes, and propose a non-microbial source for tyramine levels in white women. Finally, we demonstrate that supervised learning models trained on metabolomics data can accurately predict subsequent sPTB, potentially paving the way to new diagnostic panels.

To obtain insights into microbial metabolism of tyramine by the microbiome, we used community-scale metabolic models. These models have important limitations. Major efforts were made to curate models for gut microbes, yet such models are still missing for some vaginal microbes, and may lack representation of niche-specific metabolic capabilities. Another limitation stems from the resolution of 16S rRNA gene sequencing, which identifies taxa at the species or genus level, precluding the tailoring of models to specific strains present in each sample. Despite these limitations, our models provided accurate predictions of tyramine levels, and offered insights regarding its sources in the context of its association with preterm delivery.

We detected a group of metabolites that was largely independent from the microbiome, and which prior literature suggests to be of exogenous source, potentially from cosmetic or hygienic products. Our results coincide with recent studies which raise concern regarding environmental exposures in pregnant women^104–106^, and demarcate the presence of these chemicals in the reproductive tract. We found that in general, enrichment of sPTB associations among xenobiotics was evident mostly in Black women, potentially reflecting disparities in environmental and exogenous exposures. Such exposure patterns could differ between cohorts, and could potentially underlie the association between racial disparities in prematurity rates and racial differences in the vaginal microbiome^15,107,108^. Further study is warranted to identify the sources of these metabolites and confirm their effects on the host, microbiome, and pregnancy outcomes.

Our predictive modeling approach has several noteworthy limitations. Our use of a case-control cohort enriched for preterm deliveries limits our ability to assess population-level predictive value, and further validation is required in prospective studies. Furthermore, as our cohort was focused on sPTB, we are unable to assess if our models are specific to sPTB or are detecting a general risk for adverse pregnancy outcomes. Additionally, the untargeted panels we used do not measure metabolite concentrations, and further studies are needed for developing diagnostic panels. A larger sample size, and combination with other sources of data, such as maternal urine or serum metabolomics, vaginal metagenomics, or cell free RNA measurements, could further improve prediction accuracy. Nevertheless, our results demonstrate the potential of vaginal metabolites to serve as early biomarkers of preterm delivery and to potentially enable the personalization of treatment and prevention strategies.

## Methods

### Study design and cohort description

We analyzed banked samples from the previously collected and described Motherhood & Microbiome (M&M) cohort^15^. This cohort was approved by the Institutional Review Board at the University of Pennsylvania (IRB #818914) and the University of Maryland School of Medicine (HP-00045398), and all participants provided written informed consent. The M&M cohort recruited 2,000 women with a singleton pregnancy prior to 20 weeks of gestation. Women were followed to delivery, and spontaneous preterm birth was defined as delivery before 37 weeks of gestation with a presentation of cervical dilation and/or premature rupture of membranes. Of these, the vaginal microbiota of 503 women was characterized via 16S rRNA gene amplicon sequencing (V3-V4 region) of vaginal swabs collected between 20 to 24 weeks of gestation, and total bacterial load was assessed using the TaqMan^®^ BactQuant assay^15^. For this study, we selected, out of women with available microbiome data, all available samples from women who delivered preterm (N = 80), in addition to samples from 152 controls who delivered at term. The selected samples were replicates of those used for 16S rRNA gene sequencing, collected using a double shaft dacron swab.

### Metabolomics profiling and preprocessing

Metabolite levels were measured from vaginal swabs by Metabolon Inc. (Durham, NC, USA), using an untargeted LC/MS platform as previously described^27,109,110^. Metabolite measurements were volume normalized, followed by robust standardization^30^ of the log (base 10) transformed values (subtracting the median and dividing by the standard deviation calculated while clipping the top and bottom 5% of outliers).

### Microbiome data processing

All microbiome-based analyses were done using data previously processed with DADA2^111^ and SpeciateIT^15^, available from Table S4 of ref. 15. A single exception to this are predictive models, which were trained on 97%-clustered OTUs using the USEARCH pipeline^112^. We obtained raw sequences from the database of Genotypes and Phenotypes (dbGaP) under study accession: phs001739.v1.p1. Primers were aligned to reads and then trimmed, followed by end merging and quality filtering (-fastq_maxee 1.0). The filtered reads were then pooled together, dereplicated, clustered with a 97% threshold, and chimera filtered with the UPARSE algorithm to produce the OTU count matrix.

### Global microbiome and metabolome structure

PERMANOVA analysis was performed using Bray-Curtis distance for microbiome data and the Canberra dissimilarity metric for metabolites data. De novo clustering of metabolite vectors was done using K-medoids algorithm, also with Canberra dissimilarity. This metric is robust to outliers and sensitive to differences in common features; used with metabolomics data, it previously produced robust results under bootstrapping and generated compact clusters corresponding with prior knowledge^113^. We determined the optimal number of clusters by comparing the within cluster sum of square error and the gap statistic for clustering solutions with K between 1 and 15 (**Fig. S3**). Uniform Manifold Approximation and Projection (UMAP)^114^ was performed using the Python umap-learn package^114^, with n_neighbors=15 and min_dist=0.05 for microbiome data and n_neighbors=15 and min_dist=0.25 for metabolomics data.

### Differential abundance testing and metabolite set enrichment analysis

Differential abundance tests between metabolite levels were done using the Mann-Whitney U test for metabolites which were present in at least half of the cases. To identify functional sets of metabolites that were perturbed between sPTB and TB, we compared, for each set, the Mann-Whitney *p* values for differential abundance between PTB and sPTB for metabolites within the set to the same *p* values for metabolites outside the sets, using an additional Mann-Whitney U test. We calculated significance by comparing the *p* value of the latter test to 10,000 similar *p* values calculated on random permutations of sPTB and TB labels. For functional sets, we used definitions of super and sub pathways provided by Metabolon, as well as KEGG^56^ pathways. FDR correction was performed separately for each metabolite set type.

### Microbe-metabolite correlations

To identify associations between microbes and metabolites we estimated microbial absolute abundance by multiplying the relative abundances of each taxa by the total 16S rRNA copy number for the sample, and calculated Spearman correlations with levels of metabolites we found to be associated with sPTB across the entire cohort (**Fig. 2a)**. Spearman correlations were performed for microbe-metabolite pairs with at least 50 paired measurements, using available values without imputation, and correction for multiple testing was performed via the Benjamini-Hochberg FDR method. Edges in the microbe-metabolite network **(Fig. 3a)** were drawn for all correlations with FDR < 0.1 and absolute spearman correlation above 0.25. To determine whether edges in our network **(Fig. 3a)** were influenced by race, we compared these microbe-metabolite correlations in Black women to the same correlations in white women using a two sided Fisher R-to-z transform.

### Creating and interrogating vaginal microbiome models

Microbiome metabolic modeling was done using Microbiome Modeling Toolbox (COBRA toolbox commit: 71c117305231f77a0292856e292b95ab32040711)^90,115^, using models from AGORA^116^. All computations were performed in MATLAB version 2019a (Mathworks, Inc.), using the IBM CPLEX (IBM, Inc.) solver.

We first matched between species detected in microbiome samples and those present in AGORA^116^ (**Table S3**). Models for *Atopobium vaginae* (PB189-T1-4) and *Gardnerella vaginalis* (14019-MetR) were obtained from the AGORA authors^117^. To increase the representativeness of our models, we selected genus-level representatives for abundant vaginal species without a corresponding AGORA model that were present with >5% relative abundance in more than 20 samples: As a common *Prevotella* species in our cohort did not have a species assignment, we grouped *Prevotella* abundances to the genus level, and selected the AGORA model for *Prevotella timonensis* as its representative, as it was the most abundant taxa for which a correspondent metabolic model was present in AGORA. Similarly, we represented *Atopobium* OTUs without a species assignment with the model of *Atopobium rimae*, which was detected in vaginal samples and was the most extensive *Atopobium* model in AGORA (1054 reactions). A missing *Megasphaera* sp_2 model led us to choose *Megasphaera elsdenii* as a representative for *Megasphaera*, as this was the only AGORA model present for this genus. *Candidatus Lachnocurva vaginae* (BVAB1) was discarded, as no suitable AGORA model was available for it. To generate species level models, we combined metabolic models from available strains using the function createPanModels.m of the Microbiome Modeling Toolbox^90^. Altogether, our microbiome metabolic models included 62 different species, with an average of 15 species in each sample and a maximum of 33.

To support and improve the accuracy of our tyramine predictions, we validated the presence of the TDC gene, coding for tyrosine decarboxylase. For each species represented in our metabolic models (N = 62), we used Prodigal^126^ to predict open reading frames in up to 200 randomly selected Refseq^118^ assemblies, and searched them for evidence of TDC using the hmmsearch function of Hmmer3.3.2^119^ and a profile hmm for TDC^120^ (NCBI HMM accession: TIGR03811.1). We then curated our metabolic models, making sure that the corresponding reaction exists in models for which at least one assembly contained the corresponding gene.

For each sample, tailored microbiome models were created through the compartmentalization technique^121^: metabolic reconstructions of all species present in the sample were merged into a shared compartment, and input and output compartments were added. The shared compartment enables microbes to share metabolites while input and output compartments are present to enable compounds intake and secretion. Coupling constraints were added as in refs. 122 and 123 to ensure a dependency between relative abundances and each species network fluxes. Finally, personalized microbiome biomass objective functions composed by the sum of each microbial biomass multiplied by the corresponding relative abundance value, were added to each microbiome model.

Metabolic modeling requires environmental conditions such as media and carbon source availability^91^. We therefore formulated a “general vaginal media” (**Table S4**), as the union of all metabolites present in at least 25% of our samples to which a corresponding metabolite was identified in AGORA, assuming them to be present in an unlimited (i.e., very high) concentration. This vaginal media was applied to each microbiome model input compartment in the form of constraints on metabolite uptake reactions, constraining uptake of compounds not present in the environment to zero. To interrogate the secretion potential of each sample-specific microbiome model, we computed Net Maximal Production Capabilities (NMPCs) using the pipeline mgPipe.m of the Microbiome Modeling Toolbox^90^ (**Table S5**). We computed Spearman correlations between NMPCs and the corresponding metabolite measurements without imputation.

### Training and testing of sPTB classifiers

We constructed multiple predictive models separately using the clinical, microbiome, and metabolomics data, as well as a combination model consisting of all of these data types combined. Samples were split into training and test sets using 10-fold cross validation, block-stratified for deciles of gestational age at birth (GAB), and for microbiome, metabolomics, and combined models, also stratified for race. Train-test sterility was strictly maintained. We used LightGBM^124^ to predict GAB, and used 1,000 iterations of a random grid search to tune hyperparameters (**Table S6**). The predictions of the model were standardized for each fold separately. The final model was selected as the model with the top R^2^ score out of the top 5 most accurate models based on auROC for sPTB classification. To increase robustness, each model was evaluated on 5 randomly-selected 10 cross-validation folds. The selected models were then evaluated, without retraining, on classification of extremely (GAB < 28 weeks) or very (GAB < 32 weeks) PTB. We assessed the significance of the difference in auROC between two models by computing z-scores of the normal distributions of auROC on 30 10-fold cross-validation splits^125^.

Maternal clinical data included age, race, parity status, history of sPTB, and BMI. When training the model based on this data we calculated, for the training data of each cross-validation fold, the Spearman correlations between each feature and GAB, and then selected the top 4 features, without examining the test fold. This selected age, race, parity status, and history of sPTB in all models. The final hyperparameters of this model were learning_rate = 5e-4, num_iterations = 3500, num_leaves = 48, min_data_in_leaf = 25, max_depth = 7, feature_fraction = 0.87, and bagging_fraction = 0.64.

As race had very strong interactions with microbiome and metabolomics data, we trained a composite predictor for microbiome, metabolomics, and combination models. For microbiome data, models were trained separately for samples from Black (N = 173) and non-Black (N = 59) women. The metabolomics and combinations models followed the same scheme, but ignored samples from non-white, non-Black women (N = 8) when training. Despite the smaller sample size for each model, this empirically improved prediction performance.

Microbiome-based models used absolute abundances, calculated from USEARCH-processed OTUs as described above. For the microbiome model used for samples from Black women, the data was log-transformed and the following parameters were used with LightGBM: learning_rate = 0.01, num_iterations = 2000, num_leaves = 54, min_data_in_leaf = 20, max_depth = 9, feature_fraction = 0.96, bagging_fraction = 0.78). For the microbiome model used for samples from non-Black women, we removed the 40% most sparse features (highest fraction of zero entries) within the training data. We further performed feature selection, first by fitting a LightGBM model (with the same parameters) to the training data and then removing the bottom 10% of features based on SHAP^98^ values; and then again using Spearman correlations with GAB, removing the bottom 10% of features. The LightGBM model in this case was used with the following parameters: learning_rate = 0.05, num_iterations = 3000, num_leaves = 98, min_data_in_leaf = 19, max_depth = 8, feature_fraction = 1, bagging_fraction = 1.

Metabolomics-based models used raw metabolites data, mean-imputed and then standardized in every feature using only train data. For the model used for samples from Black women, we performed feature selection using Spearman correlation, removing the bottom 20% of features. We used the following parameters for LightGBM: learning_rate = 0.01, num_iterations = 540, num_leaves = 47, min_data_in_leaf = 24, max_depth = 5, feature_fraction = 0.95, bagging_fraction = 1. For the model used for non-Black women, we again removed the 35% most sparse features, and performed feature selection using Spearman correlation, removing the bottom 15% of features. We used the following parameters for LightGBM: learning_rate = 0.03, num_iterations = 550, num_leaves = 29, min_data_in_leaf = 1, max_depth = 4, feature_fraction = 0.95, bagging_fraction = 1.

For the combined model, we used microbiome absolute abundances. We used dimensionality reduction, which had better performance empirically, while maintaining train-test sterility, using PCA (nPC = 50) for the model used for Black women, and kernel PCA (nPC = 25) for the model used for non-Black women. The metabolomics data in the model were mean-imputed and standardized as mentioned above. We applied feature selection for the combination model used for samples from Black women, using Spearman correlation to remove the bottom 10% of features. We used the following parameters for LightGBM: learning_rate = 0.02, num_iterations = 560, num_leaves = 68, min_data_in_leaf = 24, max_depth = 7, feature_fraction = 0.95, bagging_fraction = 1. For the combination model used for non-Black women, we removed the 30% most sparse features, and then performed feature selection using Spearman correlation, removing the bottom 15% of features. We used the following parameters for LightGBM: learning_rate = 2.5e-2, num_iterations = 650, num_leaves = 13, min_data_in_leaf = 1, max_depth = 4, feature_fraction = 1, bagging_fraction = 1.

## Acknowledgments

We thank members of the Korem group, Liat Shenhav, David Zeevi, and Ronald Wapner for useful discussions, and Almut Heinken and Ines Thiele for providing metabolic models. The M&M cohort was funded by the National Institute of Nursing Research (R01NR014784). One of the datasets used was obtained from the database of Genotypes and Phenotypes (dbGaP) through dbGaP accession number phs001739.v1.p1. The current study was supported by the Center for Precision Medicine at the University of Pennsylvania, the Vagelos Award provided by Columbia University Precision Medicine Initiative, and the Program for Mathematical Genomics at Columbia University. T.K. is a CIFAR Azrieli Global Scholar in the Humans & the Microbiome Program.

## Author contributions

W.F.K, F.B and M.C.L. designed and conducted all analyses, interpreted the results, wrote the manuscript, and contributed equally to this work. K.D.G. and L.A. oversaw sample processing. H.H.L, J.L. and P.G. assisted with data analysis. J.R., M.L. and M.A.E. conceived the project, designed the study, and interpreted the results. T.K. conceived and directed the project, designed the analyses, interpreted the results, and wrote the manuscript. All authors reviewed and contributed to the manuscript.

## Supplementary figures

**Supplementary Figure 1.**
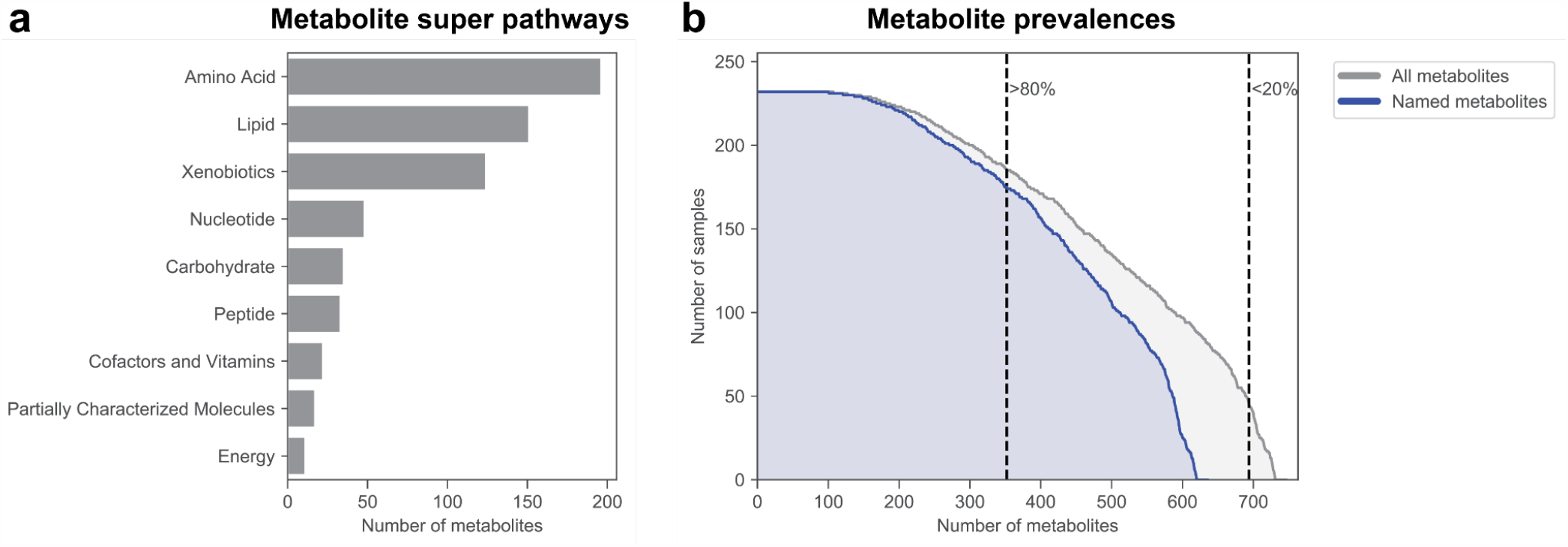
Prevalence and super pathway of assayed metabolites. **a**, Distribution of metabolite super pathways among assayed metabolites. Metabolite super pathway assignments were provided by Metabolon Inc. (Durham, NC, USA). **b**, Distribution of metabolite prevalences across samples. Gray distribution reflects prevalences of all metabolites (N = 748). Blue distribution only reflects prevalences of named metabolites (N = 637). Dashed lines distinguish metabolites prevalent in more than 80% (N = 352) and more than 20% of samples (N = 694).

**Supplementary Figure 2.**
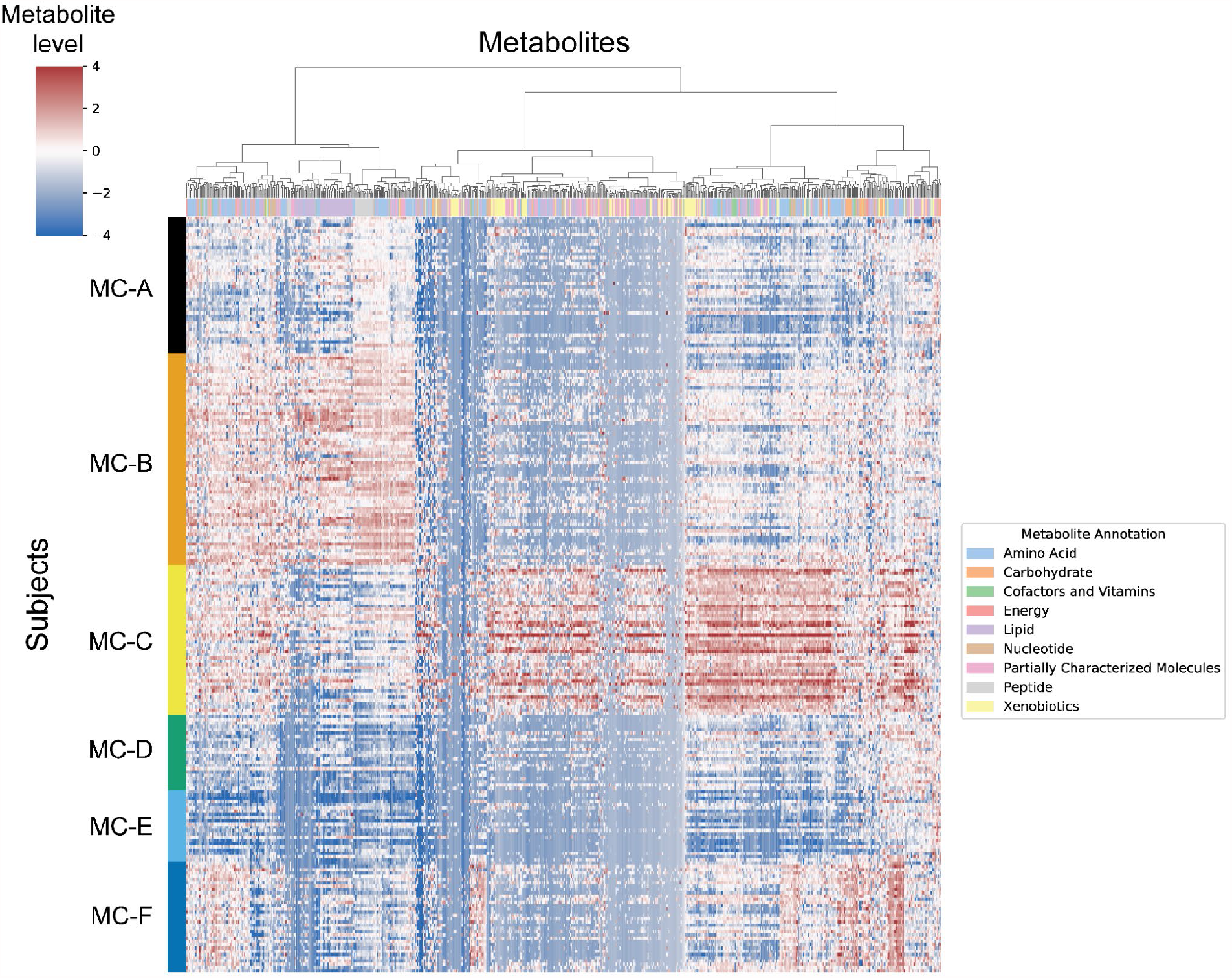
Characteristics of metabolite clusters. Heatmap showing metabolite levels for each subject (rows) and metabolite (columns). Subjects are sorted by their assigned metabolites cluster (MC) and metabolites are clustered hierarchically using Canberra distance and Ward linkage. The color above each column reflects metabolite annotations.

**Supplementary Figure 3.**
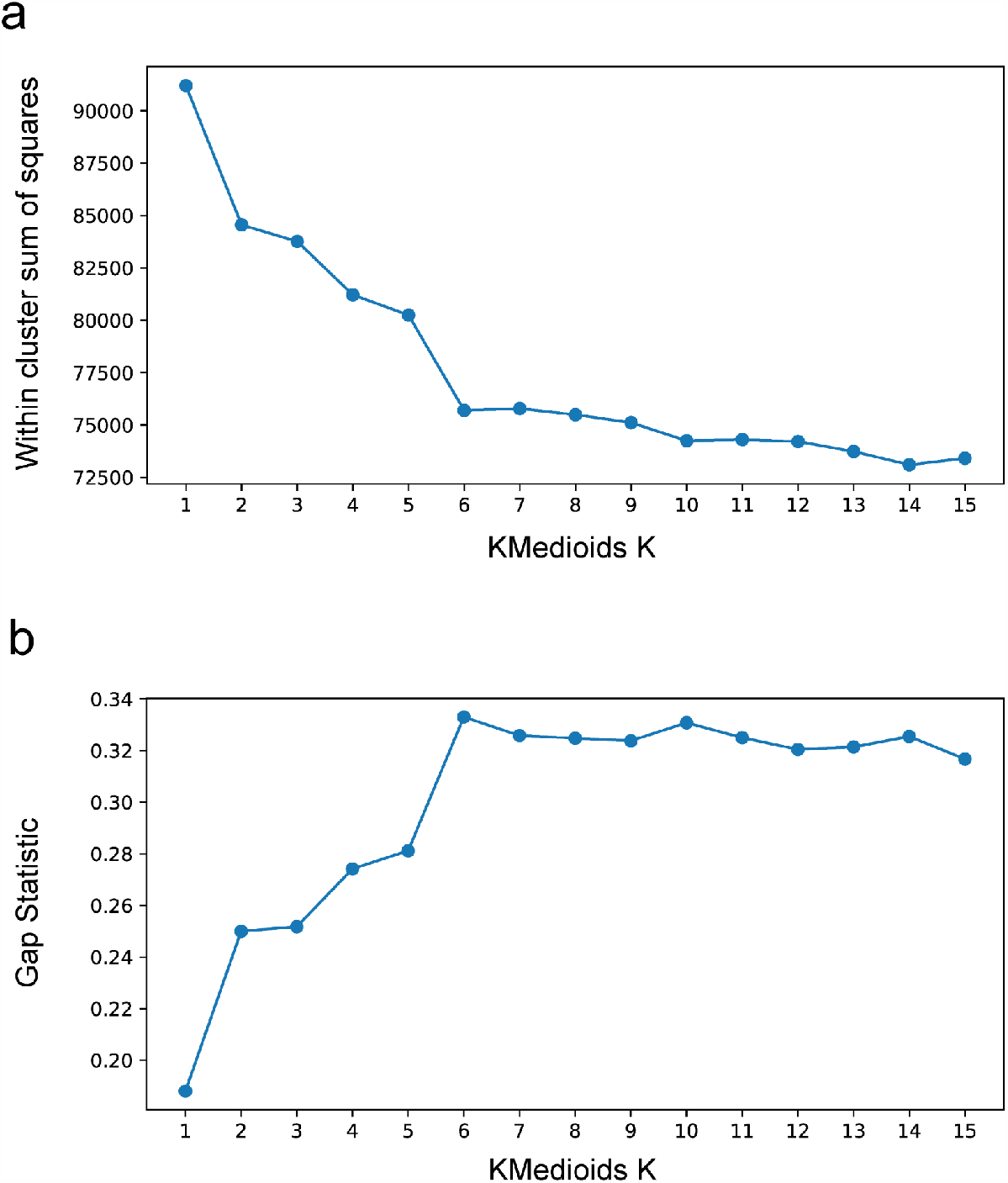
Metabolome clustering inertia and gap statistic. **a**,**b**, Within cluster sum of squared distances (a) and gap statistic (b) for k-medoids clustering using Canberra distances with k from 1 to 15. A shoulder (a) and peak (b) are visible for k = 6.

**Supplementary Figure 4.**
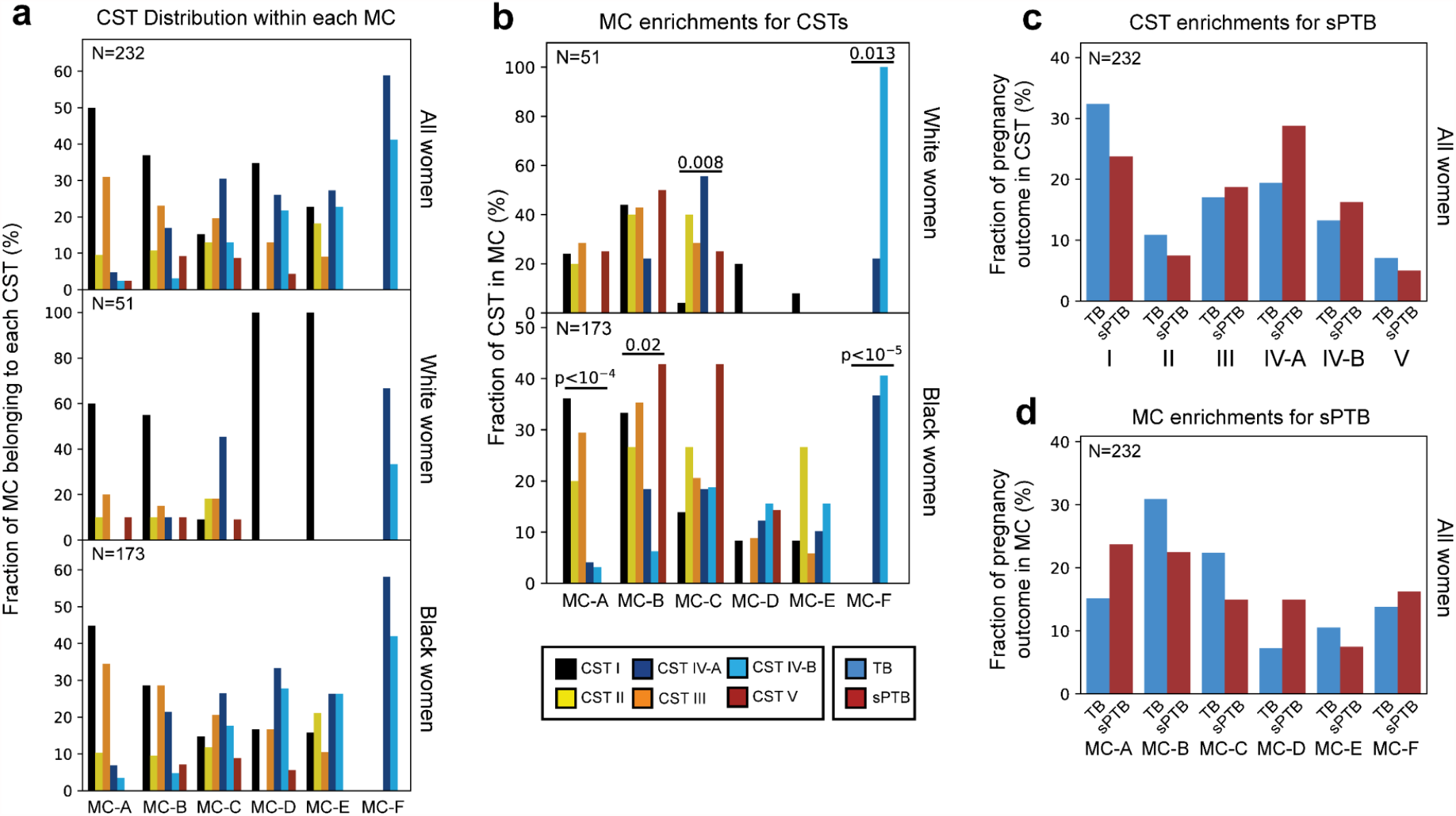
Metabolite clusters correspond to CSTs. **a**, Distribution of CSTs within each metabolite cluster, for all (top; N = 232), white (middle; N = 51) and Black (bottom; N = 173) women. Each group of bars corresponds to a single metabolite cluster and bars within a group sum to 100%. **b**, Same as **Fig. 1d**, stratified by race. **c-d**, Same as **Fig. 1f,g**, performed for all women combined.

**Supplementary Figure 5.**
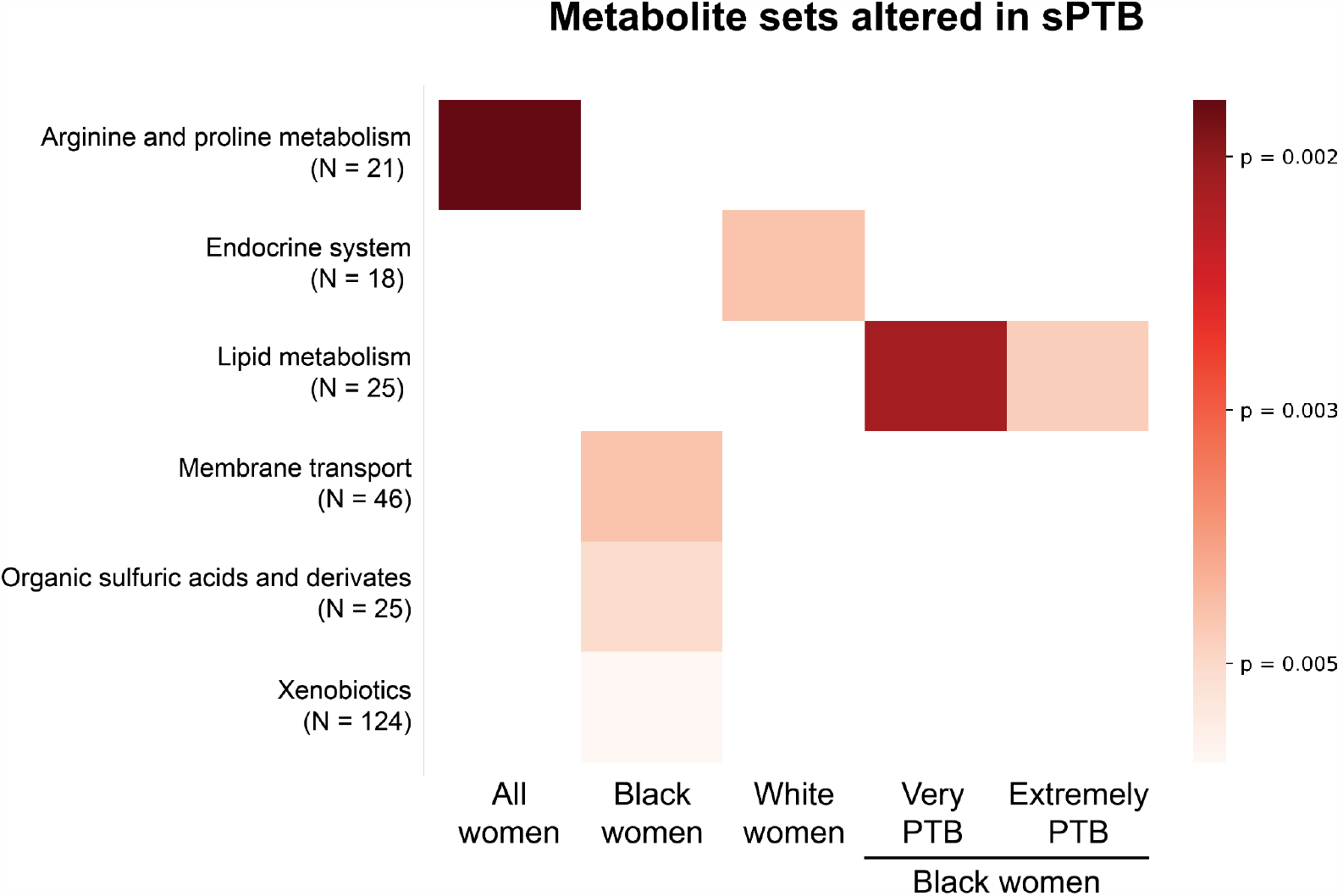
Metabolite sets altered in sPTB. Heatmap showing metabolite sets altered in sPTB in various subsets of our cohort. Colors correspond to *p*-value of metabolite set enrichment analysis (Methods). Only associations with FDR < 0.1 are shown.

**Supplementary Figure 6.**
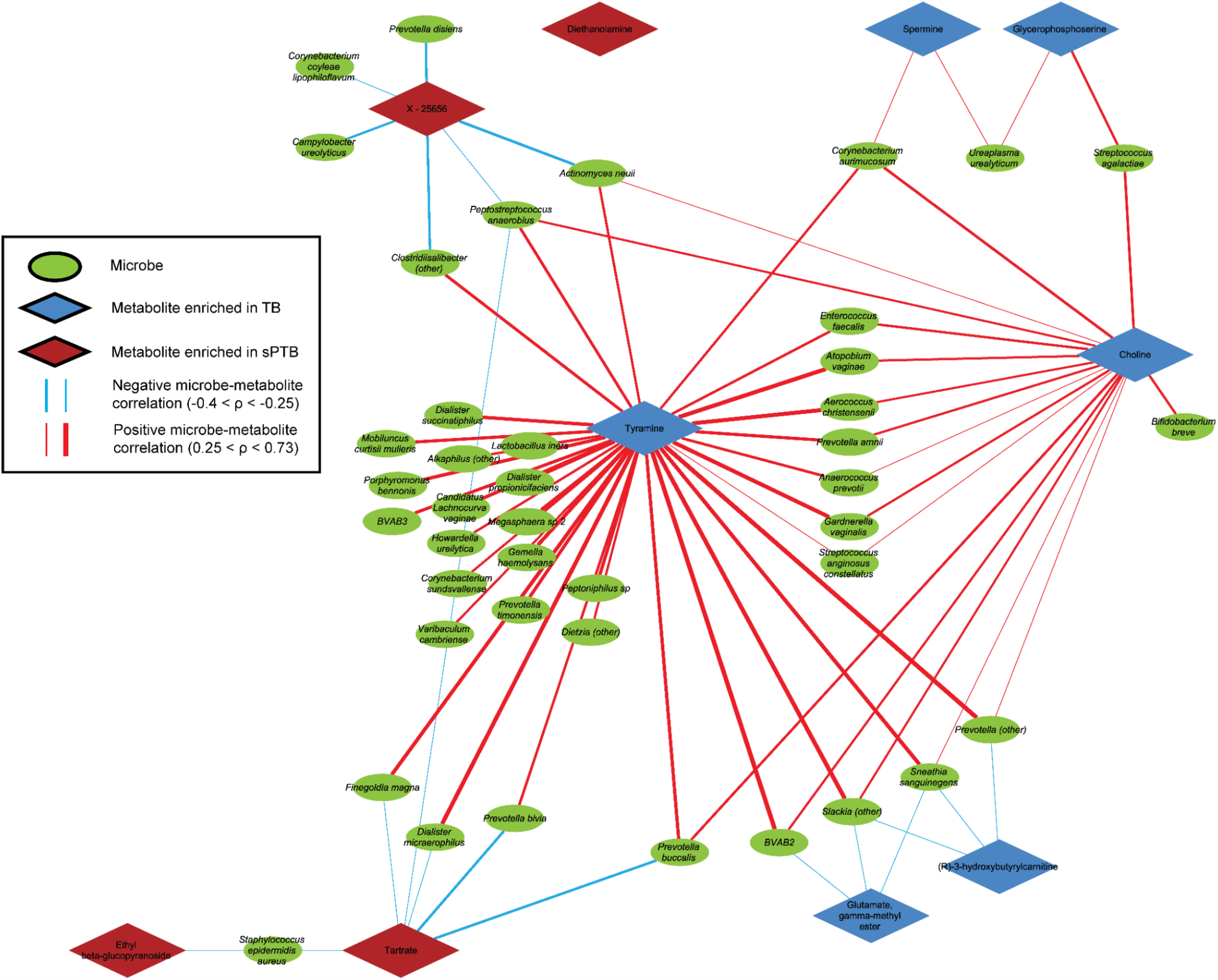
A network of microbial correlations with metabolites associated with sPTB. Same as **Fig 3a**, but with each microbial taxa represented as an individual node.

**Supplementary Figure 7.**
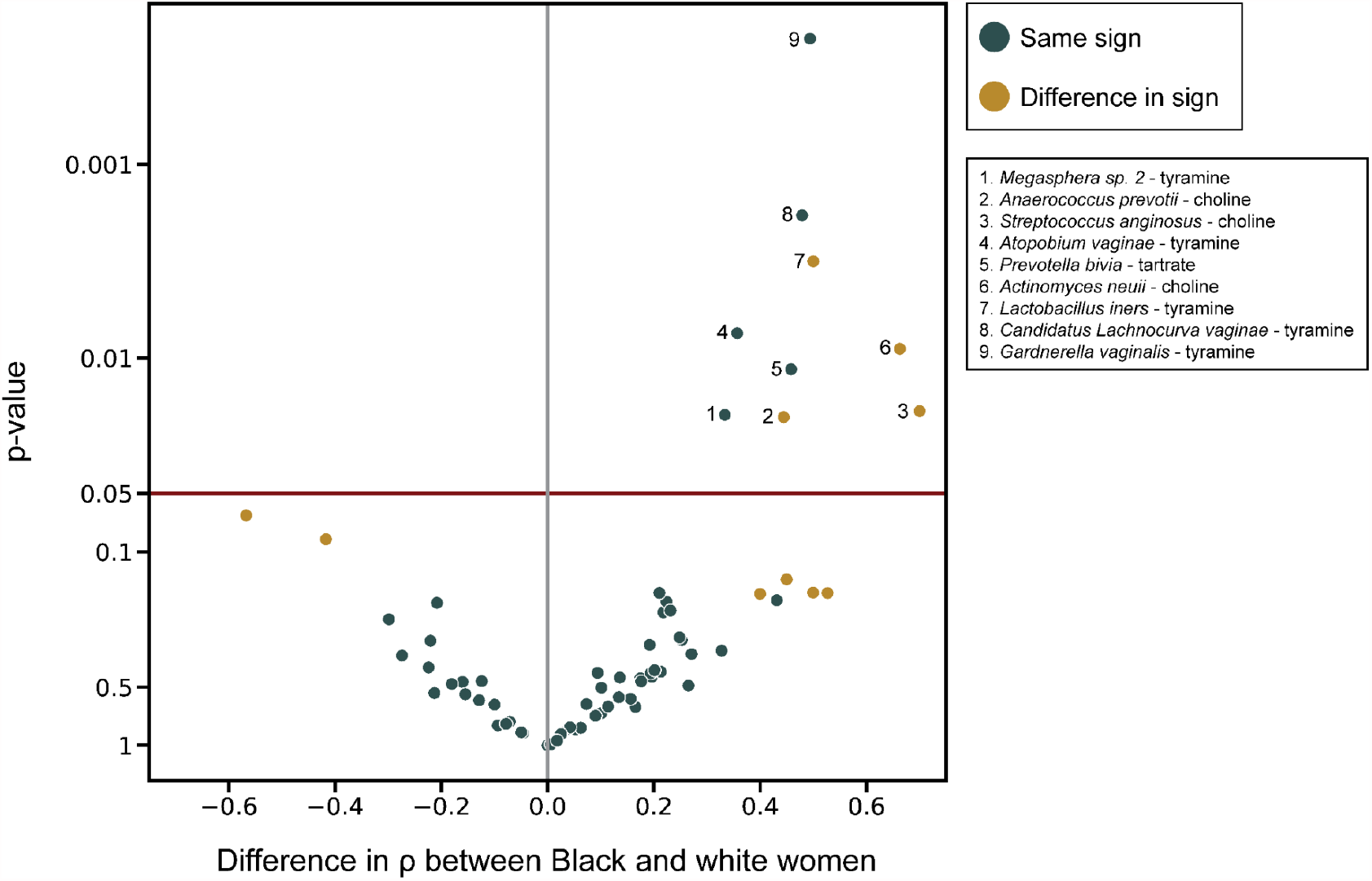
Comparison of microbe-metabolite associations in white and black women. Volcano plot where every point represents a microbe-metabolite association. X-axis displays the difference between spearman ρ’s calculated separately among Black and white women. Y-axis displays the significance of the difference, using Fisher’s R-to-z transform. Horizontal maroon line designates *p* = 0.05. Gold points indicate associations where there is a difference in sign between the correlations among Black and white women.

**Supplementary Figure 8.**
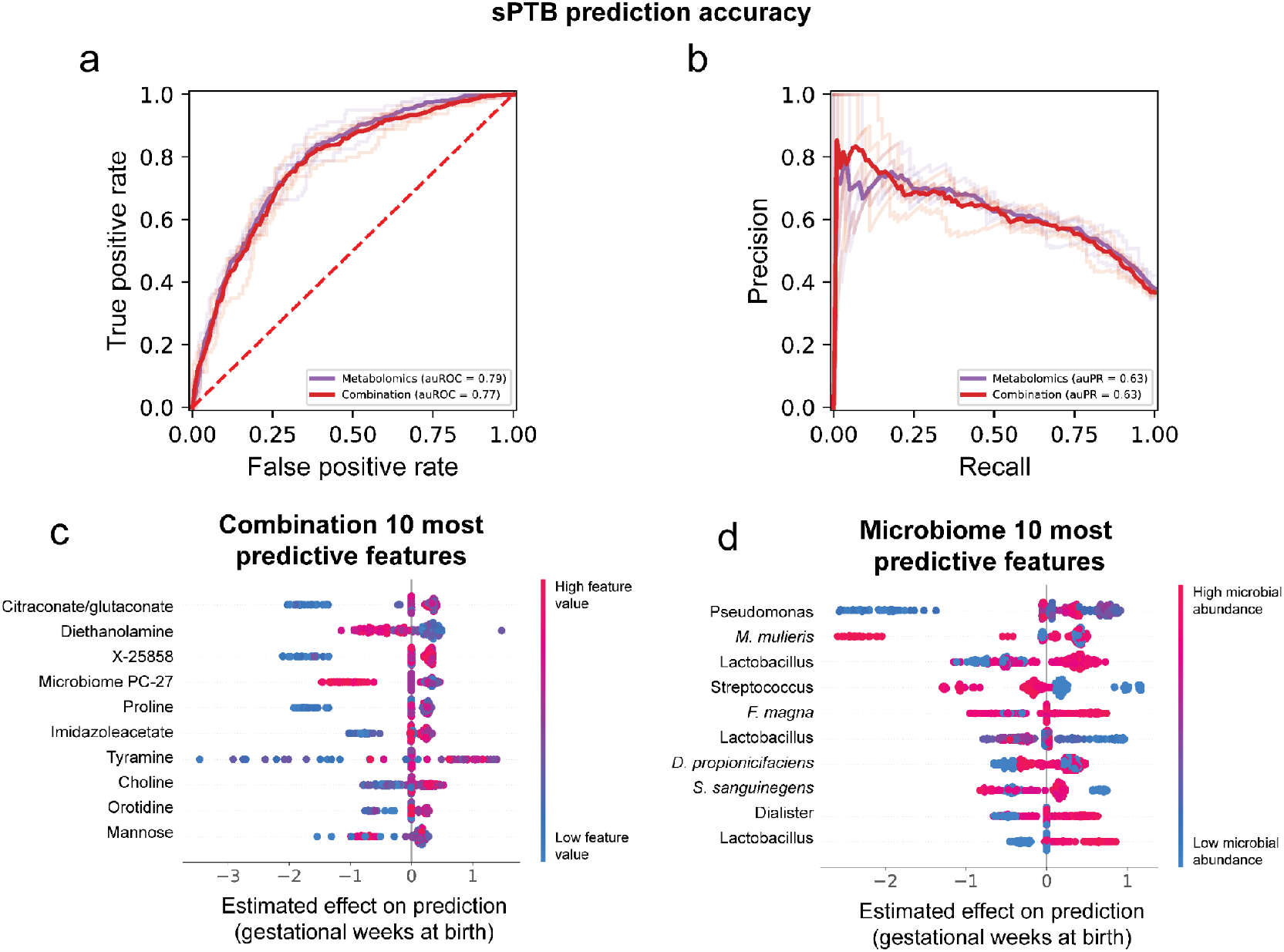
Performance and features of sPTB predictive models. **a**,**b**, Receiver operating characteristic (ROC, a) and precision-recall (PR; b) curves comparing sPTB prediction accuracy for models based on metabolomics data alone (auROC = 0.79, auPR = 0.63), and on metabolomics data combined with microbiome and clinical data (“combination”; auROC = 0.77, auPR = 0.63). N = 232 for all. **c**,**d** SHAP-based^98^ effect on total prediction (x-axis) for the top 10 features used in our combination (c) and microbiome-based (d) models, sorted with descending importance. Each dot represents a specific sample, with the color corresponding to the relative level of the metabolite, or abundance of the microbe, in the sample compared to all other samples.

## Supplementary tables

**Supplementary Table 1** | **Assignments of samples to metabolite clusters (MCs)**.

**Supplementary Table 2** | **Shapley values of prediction models**. Note that values are available only for samples on which the model was trained.

**Supplementary Table 3** | **Assignments of SpeciateIT species to AGORA models**.

**Supplementary Table 4** | **Metabolites included in the vaginal media used in metabolic models**. Listed are the metabolites included, along with their AGORA identifiers.

**Supplementary Table 5** | **Tyramine predicted Net Maximal Production Capabilities**.

**Supplementary Table 6** | **Parameters of prediction models**.

© 2021 The Trustees of Columbia University in the City of New York. This work may be reproduced and distributed for academic non-commercial purposes only without further authorization, but rightsholder otherwise reserves all rights and all reproductions of this work must include an express acknowledgment of the authors. No changes or modifications to this work may be made without prior written permission from author Tal Korem.

